# Circadian Alterations Increase with Progression in a Patient-Derived Cell Culture Model of Breast Cancer

**DOI:** 10.1101/2021.01.15.426886

**Authors:** Hui-Hsien Lin, Stephanie R. Taylor, Michelle E. Farkas

**Author notes:** **Corresponding authors:** Michelle E. Farkas, University of Massachusetts Amherst, 710 N. Pleasant St., Amherst, MA 01003, USA, Stephanie R. Taylor, Colby College, 5855 Mayflower Hill, Waterville, ME, 04901, USA.

## Abstract

Circadian rhythms are critical regulators of many physiological and behavioral functions; disruption of this time-tracking system can elicit the development of various diseases, including breast cancer. While multiple studies have used cell lines to study the correlation between altered circadian rhythms and cancer, these cells generally have different genetic backgrounds and do not mirror the changes that occur with disease development. Isogenic cell models can represent and recapitulate changes across cancer progression. Hence in the present study, a patient-derived breast cancer model, the 21T series, was used to evaluate changes to circadian oscillations of core clock protein transcription and translation as cells progress from normal to malignant states. Three cell lines from the series were used: H16N2, from normal breast epithelium; 21PT, from Atypical Ductal Hyperplasia; and 21MT, from Invasive Metastatic Carcinoma. Both of the cancerous cell lines are HER2 positive. We assessed the transcriptional profiles of two core circadian clock proteins, BMAL1 and PER2, which represent a positive and negative component of the molecular oscillator. In the normal H16N2 cells, *BMAL1* and *PER2* both possessed rhythmic mRNA oscillations with close to standard periods and the expected anti-phase relationship. However, in the cancerous cells, consistent changes were observed: both clock genes had periods that deviated farther from normal and did not have an anti-phase relationship. To provide a more complete understanding of circadian alterations in breast cancer, luciferase reporters and real-time luminometry should be used in future studies.

## Introduction

The circadian clock is a hierarchical timing system that regulates physiological, behavioral, and metabolic functions across a 24-hour day-night cycle, and maintains temporal tissue homeostasis in coordination with the external environment [1, 2]. Peripheral clocks are entrained by a central core clock located in the suprachiasmatic nucleus (SCN), and are necessary for normal tissue functioning, including cell development [3, 4]. At the molecular level, the core clock consists of a well-characterized transcriptional-translational feedback loop (TTFL), where CLOCK and BMAL1 bind to an E-box promoter to drive the expression of other clock genes, including *CRY* and *PER*. Subsequently, CRY and PER accumulate in the cytoplasm during the day to form complexes, which translocate back to the nucleus to suppress the activity of CLOCK and BMAL1 [2]. Disrupted circadian rhythms have been associated with different types of cancer, including but not limited to breast [5–8], colon [9], and prostate cancers [10]. For breast cancer in particular, epidemiological studies have shown that long-term night shift workers have a higher risk of developing the disease [11–13]. Furthermore, both *in vitro* and *in vivo* studies have indicated that mutation or dysregulation of clock genes can lead to initiation of breast tumor growth and metastases [5–8].

Among women in the United States, breast cancer is the most commonly diagnosed cancer type, with the second highest cancer death rate following lung and bronchus cancer [14]. Female breast tissue is subject to more frequent remodeling than other types [15]. For example, during pregnancy, lactation, and involution, the mammary epithelium undergoes multiple cycles of proliferation and cell death [16, 17]. As a result, the mammary gland is more prone to abnormalities, which can result in cancer [18]. While it has been shown that genetic mutations and epigenetic modifications contribute to breast cancer progression and development [19, 20], the molecular mechanisms of such processes are still unclear.

Various breast cancer cell lines (e.g. MCF-7, MDA-MB-231, SKBR3, and many others) are frequently used for studying the disease and potential therapeutics for treating it [21–23]. Although these cellular models are generally categorized into different breast cancer subtypes [24], they each represent only a single stage of a progressive disease. As a complementary approach, series of cell lines representing disease progression can be used. There are a few such models that have been used to study breast cancer development, including the MCF10 human isogenic series [25–27] and the 4T1 syngeneic mouse model [28, 29]. While each represents different stages of cancer and can be related to human disease, the MCF10 series was derived via laboratory-based genetic manipulations (including an H-Ras mutation rarely found in breast cancer patients), and the 4T1 series originated from a single, spontaneously arising mammary tumor in a BALB/c mouse. Neither may truly reflect the natural evolution of human breast cancer.

Among progressive cancer models, the 21T series of cells is particularly relevant since each cell type was isolated from the same patient, who originally had infiltrating and intraductal carcinoma, and later developed metastases to the lung [30]. Four cell lines comprise the model: H16N2, tumor adjacent non-cancerous breast cells; 21PT, Atypical Ductal Hyperplasia (ADH) cells, which are non-tumorigenic and non-metastatic; 21NT, Ductal Carcinoma In Situ (DCIS), which are tumorigenic and non-metastatic; and 21MT, Invasive Metastatic Carcinoma (IMC), which are both tumorigenic and metastatic. Two distinct populations exist within the 21MT cell line, which was further separated into 21MT-1 and 21MT-2. 21MT-1 are highly heterogeneous, tend to grow in clusters, and do not form confluent monolayers, while 21MT-2 are homogeneous polygonal cells that grow as monolayers [30]. All cancerous cells of the 21T series overexpress *ERBB2* compared to normal H16N2 cells [30]. *ERBB2* encodes the human epidermal growth factor receptor 2 (HER2), which is overexpressed, amplified, or both in several human malignancies including breast, ovarian, and colon cancers [31]. Overexpression of HER2 in human tumor cells is closely associated with increased angiogenesis, higher rates of cell survival, and poor clinical outcomes [32].

Disruption of circadian rhythms has been shown to trigger cancer development and, vice versa, malignant transformations have led to disturbances of circadian clocks [5–8, 13]. In one of our previous studies, we showed that circadian clocks are disrupted across the MCF10 series of cells representing human breast cancer [33]. We found that from benign to metastatic states, *PER2* exhibited relatively stable oscillations compared to *BMAL1*, whose periods were altered over a wide circadian range. As mentioned above, while the MCF10 series represents progressive stages of breast cancer, it originated with an H-Ras mutation, not commonly found in patients with the disease. To evaluate circadian changes in a more relevant model, we used the human 21T series of cells in the present study. This work represents the first assessment of circadian oscillations in a patient-derived model of breast cancer.

We used three cell lines from the 21T series, H16N2, 21PT, and 21MT-1, representing disease progression from benign to metastatic, to determine changes in the expression profiles of core circadian clock genes. We focused on a positive and negative component of the molecular clock oscillator, *BMAL1* and *PER2*, respectively. The mRNA expressions of *BMAL1* and *PER2* were tracked in a time-dependent manner via RT-PCR assessments. Our transcriptional data indicate that the non-oncogenic H16N2 cells possess robust circadian patterns, while 21PT and 21MT-1 have consistently less reliable rhythms, with altered characteristics. These results support our hypothesis that circadian rhythms deviate from normal when cells undergo malignant transformations. In further studies, a luciferase or fluorescent reporter should be used to track the promoter activity of clock genes and/or protein expression to generate high resolution data for a complete assessment of circadian parameters in this model.

## Materials and Methods

### Cell Culture

H16N2, 21PT, and 21MT-1 cell lines were obtained from Prof. D. Joseph Jerry (Veterinary and Animal Sciences, UMass Amherst), who received them directly from Dr. Vimla Band, who originally isolated the cells. Cells were maintained in MEM (Gibco), with 1% penicillin-streptomycin (Gibco), 1 mM HEPES (Hyclone), 1 mM sodium pyruvate (Gibco), 2 mM L-glutamine (Gibco), 10% fetal bovine serum (FBS; Corning), 15 μg/mL gentamicin (Fisher Scientific), 1 μg/mL insulin (Sigma), 12.5 ng/mL epidermal growth factor (EGF; Gibco), and 1 μg/mL hydrocortisone (Sigma). All cells were incubated at 37 °C under 5% CO_2_.

### Synchronization of Cells by Serum Shock

2 mL of cells at a density of 2 x 10^5^ cells/mL were plated in 35 mm culture dishes and incubated for approximately 24 h to reach 100% confluence. Culture media was discarded and cells were washed once with 2 mL of phosphate-buffered saline (PBS; Gibco). Cells were then starved in MEM medium without any supplements for 12 h. After starvation, cells were serum shocked using growth medium containing 50% FBS for 2 h, followed by wash with PBS and return to starvation conditions (culture in MEM medium without any supplements).

### RNA Extraction and cDNA Synthesis

Cells were collected at the first time point immediately following serum shock (T=0), and every 4 h thereafter for 48 h. Total RNA was extracted via TRIzol Reagent (Gibco) according to the manufacturer’s instructions. Briefly, 1 mL of TRIzol was added to lyse the cells, and cell lysates were transferred to microcentrifuge tubes. Cell lysates were incubated at rt for 5 min to allow complete dissociation of nucleoprotein complexes. After addition of 200 μL chloroform per 1 mL TRIzol, samples were shaken vigorously by hand for 15 s and incubated at rt for 3 min. Samples were then centrifuged at 12,000 x g for 15 min at 4 °C to separate the RNA-containing, upper aqueous phase, from the lower chloroform phase. RNA samples were further purified via PureLink RNA kit (Ambion) according to the manufacturer’s instructions. Total RNA concentration was determined via Nanodrop UV/Vis (Thermo Fisher Scientific). 1 μg of total RNA was reverse-transcribed to cDNA using 50 μM random hexamers, 40 U/μL RNaseOut, 10 mM dNTPs, and 200 U/μL SuperScript IV Reverse Transcriptase (Thermo Fisher Scientific).

### Real-Time PCR (RT-PCR)

RT-PCR was performed in 96-well plates. Each reaction (20 μL per well) consisted of 100 ng cDNA, 10 μL iTaq universal SYBR Green Supermix (Biorad), 4 μM of each respective forward and reverse primer, and RNAse free water (Fisher) to a final volume of 20 μL. All DNA primers were purchased from Integrated DNA Technologies. The following sequences were used: *GAPDH* forward (5’-CTT CTT TTG CGT CGC CAG CC-3’), reverse (5’-ATT CCG TTG ACT CCG ACC TTC-3’); *BMAL1* forward (5’-CTA CGC TAG AGG GCT TCC TG-3’), reverse (5’-CTT TTC AGG CGG TCA GCT TC-3’); *PER2* forward (5’-TGT CCC AGG TGG AGA GTG GT-3’), reverse (5’-TGT CAC CGC AGT TCA AAC GAG-3’). After brief centrifugation, samples were analyzed via CFX Connect real-time system (Biorad) programmed with an initial activation at 95 °C for 3 min, followed by 40 cycles of 95 °C denaturation for 10 s, and 60 °C annealing/extension for 30 s. Relative *BMAL1* and *PER2* expression levels were determined by comparing the *C_t_* values of *BMAL1* and *PER2* to *GAPDH* controls via the 2^ΔΔC_t_ method [34]. Three biological replicates and three technical replicates per biological replicate were analyzed for each condition.

### Rhythmicity Tests

Rhythmicity tests were performed using R packages (www.r-project.org). RAIN (rain v1.14.0) was used with a period of 24 hours. The Lomb-Scargle permutation test (randlsp from lomb package v1.2; using the peak of the periodogram in the range 6 to 50 hours) used 5,000 random permutations [35]. Metacycle v1.2.0 [36] was used to run the JTK-Cycle test. ECHO (echo.find v3.0) was used to test for periodicity in the range of 24 to 32 hours, after linearly de-trending [37].

### Curve-Fitting

Each mRNA time series (with 5-6 biological replicates) was fit to a “damped” cosine curve with a linear baseline: 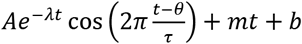, where *A* is amplitude, λ is damping rate, θ is phase, τ is intrinsic period, *m* is baseline slope, and *b* is y-intercept. λ was allowed to vary between −0.05 (a growing curve) and 0.05 (a damping curve). As in ECHO, each point was weighted by the inverse of the variance at its time step. Also, as in JTK-Cycle and ECHO, we used Kendall’s τ to measure the goodness of fit (GOF), as it is not as influenced by outliers as alternative methods (such as unity minus the mean squared error divided by the variance, often referred to as GOF). Kendall’s τ measures the fraction of pairs of points that vary concordantly, with 1 indicating perfect concordance (a form of correlation), 0 indicating a lack of correlation, and −1 indicating perfect discordance (a form of anti-correlation).

## Results

In the current study, we sought to evaluate circadian alterations in a progressive, patient-derived breast cancer model to ascertain whether changes occurred with disease status. The 21T series of cell lines have been considered as a unique experimental model of breast cancer progression since there is no other models that are originated from a single patient. We used three cell lines from the human breast cancer 21T series: H16N2, 21PT, and 21MT-1, which were isolated from a single patient and have the same genetic background, but different characteristics. H16N2 is derived from the normal epithelia, 21PT is derived from atypical ductal hyperplasia, and 21MT-1 is derived from pleural effusion of lung metastasis [30]. This is an excellent model to recapitulate the *de novo* processes arising during breast cancer evolution.

We focused on the expression of two core circadian clock components, one from the positive and the other the negative arm of the feedback loop, *BMAL1* and *PER2*, respectively. mRNA expression of both was evaluated for each of the three 21T cell lines, using RT-PCR (**Fig. 1**), with sampling once every four hours over the course of two cycles from two separate experiments. We sought to determine whether each timeseries was rhythmic and, if so, its circadian properties (such as peak time). To assess rhythmicity, we chose multiple, complementary methods that allowed for replicates and were flexible with respect to waveform and missing data: RAIN (arbitrary wave form) [38]; a Lomb-Scargle permutation test (allows for missing data) [35]; JTK-Cycle (compares reference curves to data) [39]; and ECHO (fits data to a sine curve with amplitude changes over time) [37]. To assess circadian properties and the quality of rhythmicity, we fit each time-series (with replicates) to a damped cosine curve. To be consistent with two of the rhythmicity tests we employed (JTK and ECHO), we determined goodness of fit using Kendall’s tau, which measures the rank correlation between the data and the fit.

**Figure 1.**
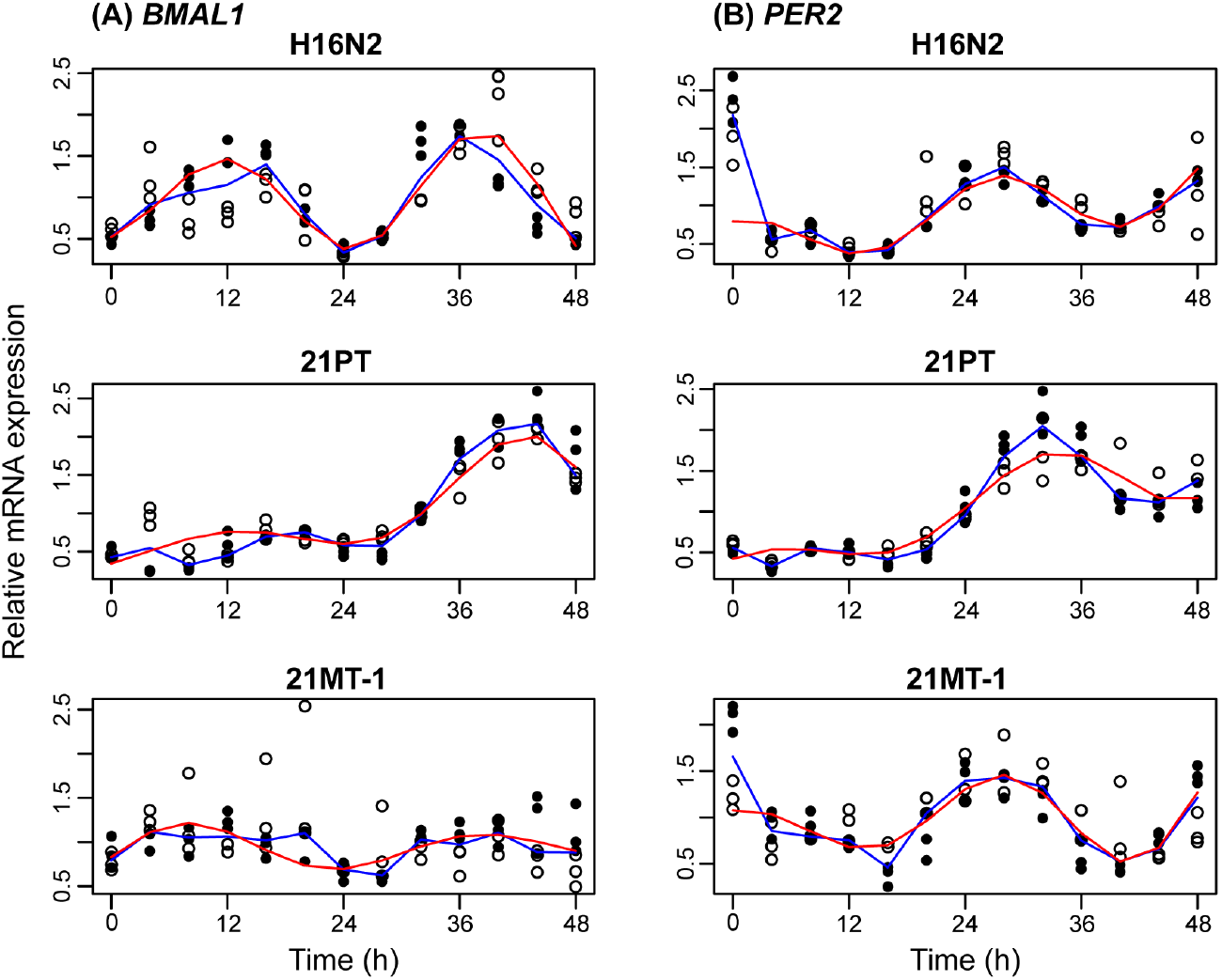
Relative mRNA expression of *BMAL1* and *PER2* across the 21T series of cells. Shown are the mRNA expression levels relative to the mean over time for each biological replicate (N=6 for all but 4 in 21MT-1, where N=5). The experiment was conducted twice (N=3 for each); open and closed circles are used to differentiate between the two. The median and best-fit damped cosine curves are shown in blue and red, respectively. Kendall’s τ identifies a low-quality fit for *BMAL1* 21MT-1 (τ=0.29) and medium-quality fits for the remaining (τ= 0.62 for *BMAL1* H16N2, 0.58 for *PER2* H16N2, 0.63 for *BMAL1* 21PT, 0.66 for *PER2* 21PT, and 0.55 for *PER2* 21MT-1).

All tests for rhythmicity indicated that the non-oncogenic H16N2 cells are rhythmic, where p < 0.001 for each with the exception of ECHO for *PER2*, which was p < 0.01 (**Fig. 2**, **Table S1** and **S2**). For 21PT, while the data for *BMAL1* are rhythmic (p < 0.001 for all tests), analyses yielded mixed outcomes for *PER2*, with p < 0.001 for the LSR and ECHO tests while p > 0.2 for RAIN and JTK) (**Fig. 2**, **Table S1** and **S2**). The patterns for the *BMAL1* and *PER2* transcripts from 21MT-1 cells were found to be rhythmic using most tests (p < 0.01; **Table S1** and **S2**), with the exception of the Lomb-Scargle permutation test for *BMAL1*, where p=0.152. Given that each data set was determined to be rhythmic by more than one test, we sought to further assess the characteristics of the oscillation patterns.

**Figure 2.**
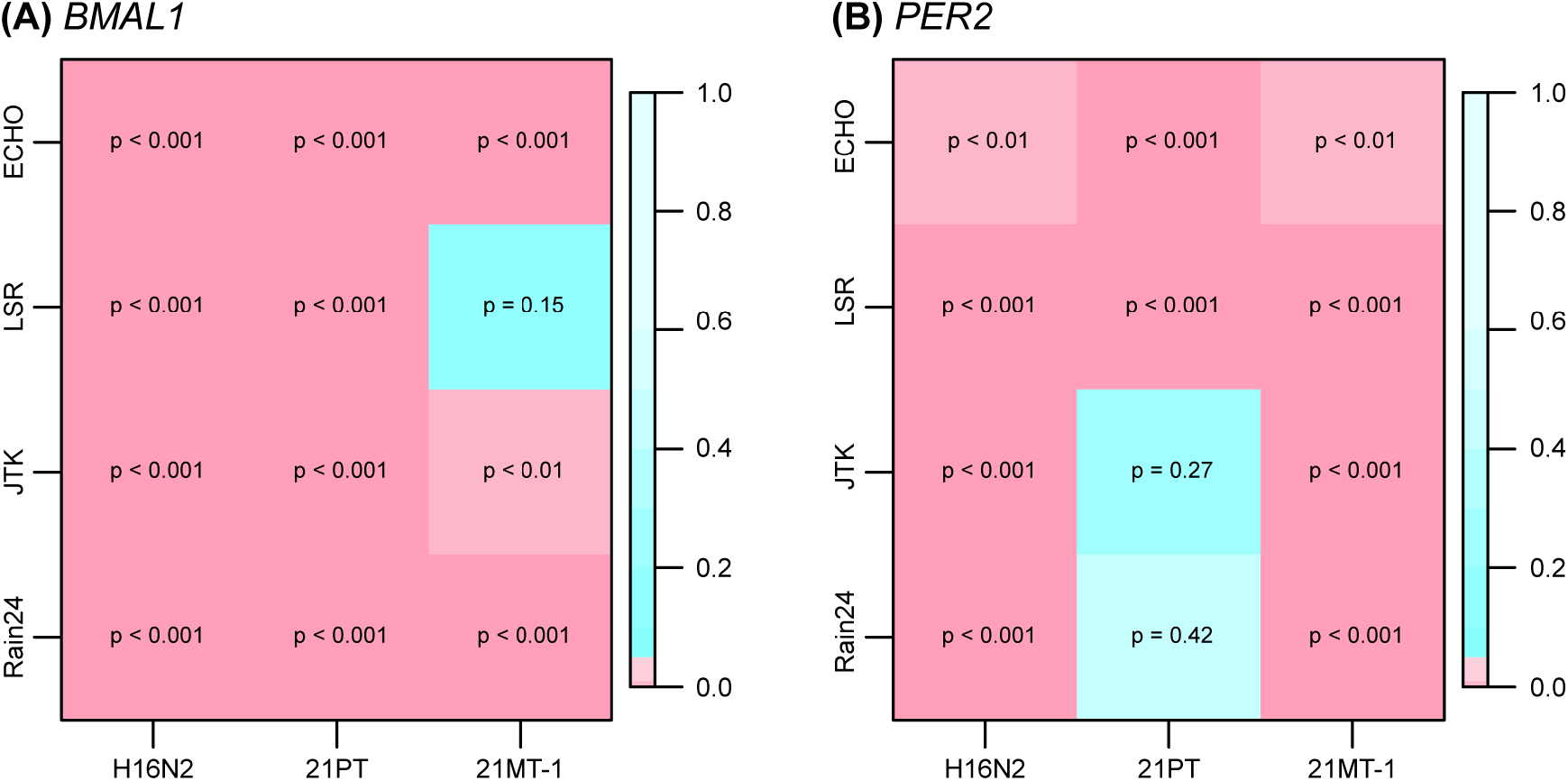
P-values determined for each rhythmicity test, cell line, and transcript. Shown is a heat map for each gene indicating p-values from 4 tests for rhythmicity applied to each cell line. Values are both labeled and color-coded (shades of blue for p ≥ 0.05, shades of pink for p < 0.05). Results indicate that H16N2 (non-cancerous) is the most consistently rhythmic across analyses accounting for both transcripts. The 21MT-1 and 21PT cell lines both include rhythmic and arrhythmic outcomes, depending on test and transcript. Rain24 = RAIN with a period of 24 h; JTK = JTK-Cycle; LSR = Lomb-Scargle Permutation test. All tests were applied to time-series with N=6 replicates per time point with the exception of 21MT-1, where N=5 for 4 time points. Exact values for each and additional tests are shown in **Table S1** and **S2**.

The normal breast epithelial H16N2 cells showed *BMAL1* peaks at ~12-16 h and 32-36 h, while those for *PER2* were at ~0-4 h and ~24-28 h. Hence, as expected for these positive and negative elements of the core clock, the phase difference between the two is approximately 12 h, with oscillation patterns in opposition to each other. A slowly amplifying cosine curve with growth rates of 0.02/h and 0.01/h for *BMAL1* and *PER2*, respectively, fit the data relatively well (Kendall’s τ=0.62 for *BMAL1* and 0.58 for *PER2*, where τ=1 indicates perfect rank correlation). It also gave more precise measures of peak times, where *BMAL1* peaked 11.4 h and *PER2* peaked 0.5 h into each cycle. The periods of both *BMAL1* and *PER2* were determined to be 26.5 h.

The 21PT (non-tumorigenic and non-metastatic) cells showed oscillations with *BMAL1* peaks at ~16-20 h and 40-44 h, while *PER2* peaks occurred at ~8-12 h and 28-32 h. Unlike in the normal cells, the transcriptional patterns here were not anti-phase but had consistent changes (an approximately 8 h offset) for both *BMAL1* and *PER2* in all samples. Curve-fitting also yielded reasonable fits for this cell line, where Kendall’s τ=0.63 for *BMAL1* and 0.66 for *PER2*. Compared to H16N2 cells, 21PT oscillations showed a larger growth rate (0.03/h for *BMAL1* and 0.05/h for *PER2)*. The periods were also both longer than those of H16N2 transcripts, and less consistent with one another, with that of *BMAL1* being 32.0 h and *PER2* 30.5 h.

The 21MT-1 (tumorigenic, metastatic) cells showed oscillations with the least circadian characteristics. The transcriptional patterns had the worst fits, where Kendall’s τ=0.29 for *BMAL1* and 0.55 for *PER2*. As in the 21PT cells, there were only 8 hours between the peaks of *BMAL1* (8 h into each cycle) and *PER2* (0 h into each cycle), but the metastatic cells showed the greatest difference in period estimates between the genes, where *BMAL1* was 30.5 h, while *PER2* was 27.1 h. The growth rates were also contrasting, with *BMAL1* diminishing at −0.03/h and *PER2* increasing at 0.05/h.

To verify the assessment that the 21PT data was the most sinusoidal and that of 21MT-1 was the least, we computed two additional measures of fit: (1) NRMSE (the root mean squared error), normalized by the data range and (2) GOF, as described in the methods section. For each period in the range of 24 to 32 h (with a step-size of 0.5 h), we fit the data to a damped cosine curve and computed the three measures of fit, including Kendall’s τ. The results were consistent across periods and fitness measures – the 21PT data fit best and 21MT-1 data fit worst, with very few exceptions (**Fig. S1**). Quality of fit for H16N2 recordings varied by measure.

Additionally, to assess possible outcomes of mRNA expression changes at the protein level, we conducted time-course western blot analyses for BMAL1 and PER2 with the three members of the 21T series of cells evaluated above (**Fig. S2**). We fit each of the time-series to damped cosine curves (**Fig. S3**). Unlike for mRNA expression patterns, the sinusoidal curves could not be adequately fit to the time-series, resulting in low goodness-of-fit measures. BMAL1 and PER2 were not observed to have a consistent phase relationship. While some of statistical tests for rhythmicity identified particular time-series as rhythmic (**Fig. S4**), these results were too inconsistent to be considered reliable, as confirmed by visual inspection.

## Discussion

Various experimental platforms have been used to investigate the correlation between altered circadian rhythms and cancer development. These include murine models with spontaneously-developing or implanted tumors [40, 41], individual cancer cell lines representing specific subtypes (e.g. MDA-MB-231 or MCF7, as mentioned above) [8, 42], and series of cells representing cancer progression (e.g. 4T1 or MCF10 series, as mentioned above) [33, 43]. Studies using these models typically show that the expression of clock genes and/or proteins are disrupted (i.e. with arrhythmic oscillation patterns or diminished expression levels) via RT-PCR or western blotting assessments. However, the data obtained may not reflect progressive oncogenic changes, since non-related cell lines have diverse genetic backgrounds and the other models were not directly derived from patients carrying breast cancer. At the same time, assessments in clinical studies often involve only a single time point [6, 7, 44, 45]. However, the nature of circadian rhythms is dynamic, and it is important to carry out direct comparisons as a function of time. We recognize the difficulty in obtaining/using patient samples in such a manner, and so, in the present study, we used primary and metastatic tumor cells along with those of the normal breast epithelium, derived from the same individual to directly compare circadian properties in a time-dependent manner.

The foundation of circadian rhythmicity is the oscillation of molecular clocks with a period of approximately 24 hours [46, 47]. During 24-hour cycling, the peak times of positive and negative core clock regulators (e.g. *BMAL1* and *PER2*, respectively) exhibit a nearly 12-hour delay from one another, resulting in an anti-phase relationship [46]. While this is the case in healthy cells, in disease states, various alterations (e.g. changes to period [47], phase [48], and amplitude [49]) may occur. As shown in **Fig. 1** and **2**, *BMAL1* and *PER2* transcripts from normal breast epithelial H16N2 cells showed rhythmic, anti-phase mRNA oscillations (~12 h differences in peak times) with similar periods that were close to 24 h (estimates were 26.5 h for both), and good fits. However, longer and dissimilar periods (32.0 h for *BMAL1* and 30.5 h for *PER2* in 21PT; 30.5 h for *BMAL1* and 27.1 h for *PER2* in 21MT-1) and altered phase relationships (~ 8 h shifts in peak times between *BMAL1* and *PER2)* were found in the cancerous 21PT (ADH) and 21MT-1 (IMC) cells.

Few studies have analyzed changes in circadian characteristics using progressive cancer models, and this is the first such study in a disease-relevant model of breast cancer. Nonetheless, we compared our findings to those of others, and strikingly observed similarities. For example, Relógio et al. discovered that metastatic human skin and colorectal cancer cell lines tended to have longer periods and altered phases compared to the respective normal epithelial cells [47, 50]. Deviations from period and phase are also seen in our data, where 21PT cells exhibited consistent changes (i.e. ~8 h phase offset with similar growth rates) for *BMAL1* and *PER2*, while 21MT-1 showed contrasting alterations for the two genes (i.e. ~8 h phase offset with two opposing growth trends, one increasing and the other decreasing), and the worst sinusoidal curve fits overall. It is surprising to us that different cancers can have similar changes in circadian characteristics. Broadly, our results support the hypothesis that circadian rhythms are increasingly disrupted in breast cancer.

Although it is unclear how circadian changes occur in the cancerous 21PT and 21MT-1 cells, molecular profiling of the cells may provide some clues. Several studies have characterized the genetic profiles of cells from the 21T series [30, 51–54]. Band et al. showed that compared to normal epithelial H16N2 cells, the others were *HER2* positive (overexpressing *ERBB2*) [30]. It is known that *ERBB2* is closely associated with *NR1D1* [55], the gene that encodes REV-ERBα (a clock protein involved in the secondary clock TTFL), and high levels of *NR1D1* have frequently been found in breast cancer patients [56]. In addition to being *HER2* positive, 21MT-1 cells had hyper-activated p-Ser^473^ Akt [52], a kinase involved in the PI3K/Akt/mTOR pathway frequently activated in breast cancer [57, 58]. AKT phosphorylates CLOCK and BMAL1, and inhibits their nuclear localization [59, 60], and the mTOR pathway has been shown to regulate circadian entrainment in the SCN [61]. Taken together, abnormal *ERBB2/NR1D1* expression and activated Akt/mTOR pathways may all contribute to the prolonged periods and altered phases of *BMAL1* and *PER2* in the cancerous 21PT and 21MT-1 cells.

To evaluate whether changes in transcriptional expression can be detected at the translational level, we performed time-course western blot analyses of BMAL1 and PER2 in the three cell lines from the 21T series. Our results showed that there was no clear and consistent phase relationship between BMAL1 and PER2 in any of them (**Fig. S2**, **S3**, and **S4**). Although some statistical analyses revealed rhythmicity in certain expression patterns (**Table S3** and **S4**), it is likely that the changes were too small to be reliable. Several studies have investigated temporal changes in clock protein expression using different disease models, however, while some showed clear and detectable protein oscillations [62, 63], others did not, perhaps due to the low resolution of western blotting [64, 65]. We also noted subtle changes in GAPDH expression over time, however, the minor instability of a reference protein has been shown previously in various cell models [50, 66, 67].

The poor correlation between observed mRNA and protein expression could also be induced by changes to transcriptional regulation, post-transcriptional modifications, protein half-lives, and technical errors and noise in either or both mRNA and protein experiments [68–70]. As an example, Robles et al. showed that post-transcriptional modifications caused a phase delay of 6 hours between cycling transcripts and corresponding proteins [69]. Future studies with this system should use luciferase reporters to acquire high-resolution data at both the promoter and translational levels, which will enable more detailed visualizations of oscillations and thorough analyses.

## Conclusion

In this study, we report the first findings of circadian disruption in a progressive series of human, breast cancer patient-derived cells. We evaluated the oscillations of mRNA expression for two core clock components, *BMAL1* and *PER2*, in a time-dependent manner, in cell lines across the 21T series. Normal epithelial H16N2 cells displayed standard oscillation patterns. The two cancerous cell types, 21PT and 21MT-1, exhibited periods that deviated farther from 24 hours and were less consistent between the *BMAL1* and *PER2* transcripts, which were also not anti-phase with the expected 12 hours difference in peak times. 21PT cells showed consistent changes in the phase relationship, with similar, growing trends for both *BMAL1* and *PER2*, while the 21MT-1 cellular transcripts were found to be less sinusoidal and had contradictory growth rates (one increasing and the other decreasing), as evidenced by diminished fits.

Our analyses of protein expression revealed that neither BMAL1 nor PER2 had clear, anti-phase relationships in any of the cells. This is likely due to the low-resolution feature of western blotting, where changes were too subtle to be detected. Taken together, our data support the hypothesis that circadian rhythms are increasingly disrupted with malignancy in breast cancer. To provide a better understanding and more accurate analyses of these altered oscillations, luciferase reporters and real-time luminometry should be used in the future. Furthermore, next generation sequencing can be utilized to assess connections between cancer and circadian pathways, and may provide insights to how they affect one another. Understanding the contributions of circadian rhythms to and their effects on breast cancer development will be pivotal to understanding roles of the clock in this and other diseases.

## Acknowledgements

H.-H. L. was supported by a University of Massachusetts Amherst Chemistry-Biology Interface (CBI) training fellowship. We are grateful to the laboratory of D. Joseph Jerry (Veterinary and Animal Sciences, UMass Amherst) for providing cell lines and culture protocols. We would like to thank Tanya Leise (Mathematics & Statistics, Amherst College), Jennifer Hurley (Biological Sciences, Rensselaer Polytechnic Institute), and Hannah De Los Santos (Computer Science) for helpful discussions regarding analyses. We are also grateful to the laboratory of Elizabeth Vierling (Biochemistry and Molecular Biology, UMass Amherst) for G:Box iChemi XT imaging system (GeneSys) access.

