## Supporting Information for "Circadian Alterations Increase with Progression in a Patient-Derived Cell Culture Model of Breast Cancer"

### Supplemental Methods

### Rhythmicity Tests

We examined both transcript and Western blot time-series with additional tests for rhythmicity. Our goal was to ensure our that results were robust to the choice of test and to better understand the limitations of commonly used tests. These included RAIN at additional periods, Harmonic Regression assessments, and Lomb-Scargle as implemented in MetaCycle. As RAIN is a method that tests for rhythmicity at a given period, in addition to the results for a test period of 24 h presented in the main text, here, we add results for periods of 20 h and 28 h. Harmonic Regression (Westermark, R package HarmonicRegression) fits the data to the first harmonics of a Fourier expansion and performs an F-test against the null hypothesis of Gaussian noise. Because it fits to a given period, we test a range of periods here, from 20 h to 28 h. It does not allow for amplitude growth or decay over time. Like the Lomb-Scargle permutation test, Lomb-Scargle as implemented in MetaCycle tests whether the peak of the periodogram is due to random noise. However, unlike the permutation test, it assumes a particular null distribution of the test statistic.

### Protein Isolation

Following TRIzol treatment and removal of the RNA-containing aqueous phase as described above, any residual aqueous phase and the interface were completely removed from the sample. 0.3 mL of 100% ethanol was added to the remaining organic phase for initial homogenization. Microcentrifuge tubes were inverted several times for mixing and incubated at rt for 3 min. The samples were then centrifuged at 2000 x g for 5 min at 4 °C to pellet any residual DNA. The phenol-ethanol supernatant, which contained protein, was transferred to a new microcentrifuge tube for further protein precipitation. An excess of isopropanol was added (at minimum 2X volume of the phenol-ethanol solution) to the phenol-ethanol phase, and the samples were incubated for 10 min at rt. Following incubation, the samples were centrifuged at 12,000 x g for 10 min at 4 °C to pellet proteins. The supernatants were then removed and discarded. 500 µL of 95% ethanol was added and microcentrifuge tubes inverted to wash the protein pellets. The samples were then centrifuged at 7,600 x g for 5 min at 4 °C. The supernatants were discarded, and an additional wash with 250 µL of 95% ethanol was performed. After decanting the supernatant, the pellets were air-dried for 30 min at rt. 100 µL of the optimized lysis buffer (adapted from Kopec et al. [1]; 20 mM EDTA, 140 mM NaCl, 5% SDS, 100 mM Tris, and 1% Halt™ Protease and Phosphatase Inhibitor (Thermo Fisher Scientific)) was added to each microcentrifuge tube, which was incubated for 45 min at 50 °C while shaking at 450 rpm. After incubation, all protein pellets were completely dissolved and protein concentrations were measured using a BCA assay (Fisher Scientific). All protein solutions were stored at -20 °C for further characterization.

**Western Blotting**

20 μg of protein per sample was electrophoretically separated on an 8% SDS polyacrylamide gel, and transferred to a PVDF membrane (Thermo Fisher Scientific). The membrane was blocked with 5% (w/v) bovine serum albumin (BSA; Fisher Scientific) in 1X TBST (150 mM NaCl, 20 mM Tris-HCl, 0.1% Tween 20) for 2 h at rt, incubated with primary antibodies against BMAL1 (Cell Signaling), PER2, and GAPDH (Proteintech) overnight at 4 °C, washed three times with 1X TBST, and incubated with horseradish peroxidase-conjugated goat-anti-rabbit IgG secondary antibody (Thermo Fisher) for 2 h at rt. Immunoblots were imaged using an enhanced chemiluminescence reagent (ECL; Thermo Fisher) via a G:Box iChemi XT imaging system (GeneSys). Band intensities were analyzed via ImageJ.

**Table S1.** Exact P-values from rhythmicity tests for the *BMAL1* mRNA time-series.

| | Rain20 | Rain24 | Rain28 | H20 | H22 | H24 | H26 | H28 | JTK | LS | LSR | ECHO |
|---|---|---|---|---|---|---|---|---|---|---|---|---|
| **N** | 6.44e-09 | 9.16e-20 | 2.56e-16 | 2.20e-07 | 6.10e-12 | 8.37e-14 | 1.07e-13 | 5.21e-12 | 1.79e-14 | 4.90e-08 | 0.00e+0 | 1.97e-16 |
| **PT** | 0.149 | 2.42e-06 | 1.94e-07 | 0.201 | 2.89e-03 | 3.05e-05 | 1.12e-05 | 3.54e-05 | 2.50e-04 | 3.63e-03 | 0.00e+0 | 1.97e-16 |
| **MT** | 0.258 | 2.71e-04 | 2.79e-05 | 0.306 | 0.106 | 3.96e-02 | 2.01e-02 | 1.48e-02 | 3.94e-03 | 0.799 | 0.152 | 4.60e-12 |

For each cell line (N=H16N2, PT=21PT, MT=21MT-1), time-series with 6 replicates per time point with the exception of 21MT-1 (which has 5 replicates for 4 of the time points), were assessed for rhythmicity with each of 12 tests. Rain20, Rain24, and Rain28 indicate RAIN with test periods of 20 h, 24 h, and 28 h, respectively; H20, H22, H26, and H28 indicate Harmonic Regression test with periods of 20 h, 22 h, 24 h, 26 h, and 28 h; JTK indicates JTK-Cycle; LS indicates Lomb-Scargle as implemented by MetaCycle; LSR indicates the Lomb-Scargle Permutation test.

**Table S2.** Exact P-values from rhythmicity tests for the *PER2* mRNA time-series.

| | Rain20 | Rain24 | Rain28 | H20 | H22 | H24 | H26 | H28 | JTK | LS | LSR | ECHO |
|---|---|---|---|---|---|---|---|---|---|---|---|---|
| **N** | 7.04e-03 | 4.92e-14 | 6.52e-18 | 0.056 | 1.10e-05 | 1.38e-09 | 1.39e-12 | 2.59e-13 | 2.41e-11 | 9.50e-08 | 0.00e+0 | 1.10e-03 |
| **PT** | 4.71e-02 | 0.424 | 0.294 | 1.58e-02 | 4.96e-02 | 0.145 | 0.220 | 0.155 | 0.273 | 0.826 | 0.00e+0 | 3.10e-13 |
| **MT** | 0.204 | 2.60e-08 | 2.95e-08 | 0.120 | 1.52e-04 | 3.43e-08 | 4.73e-11 | 3.02e-12 | 1.63e-11 | 4.64e-07 | 0.00e+0 | 1.10e-03 |

For each cell line (N=H16N2, PT=21PT, MT=21MT-1), time-series with 6 replicates per time point with the exception of 21MT-1 (which has 5 replicates for 4 of the time points), were assessed for rhythmicity with each of 12 tests. Rain20, Rain24, and Rain28 indicate RAIN with test periods of 20 h, 24 h, and 28 h, respectively; H20, H22, H26, and H28 indicate Harmonic Regression test with periods of 20 h, 22 h, 24 h, 26 h, and 28 h; JTK indicates JTK-Cycle; LS indicates Lomb-Scargle as implemented by MetaCycle; LSR indicates the Lomb-Scargle Permutation test.

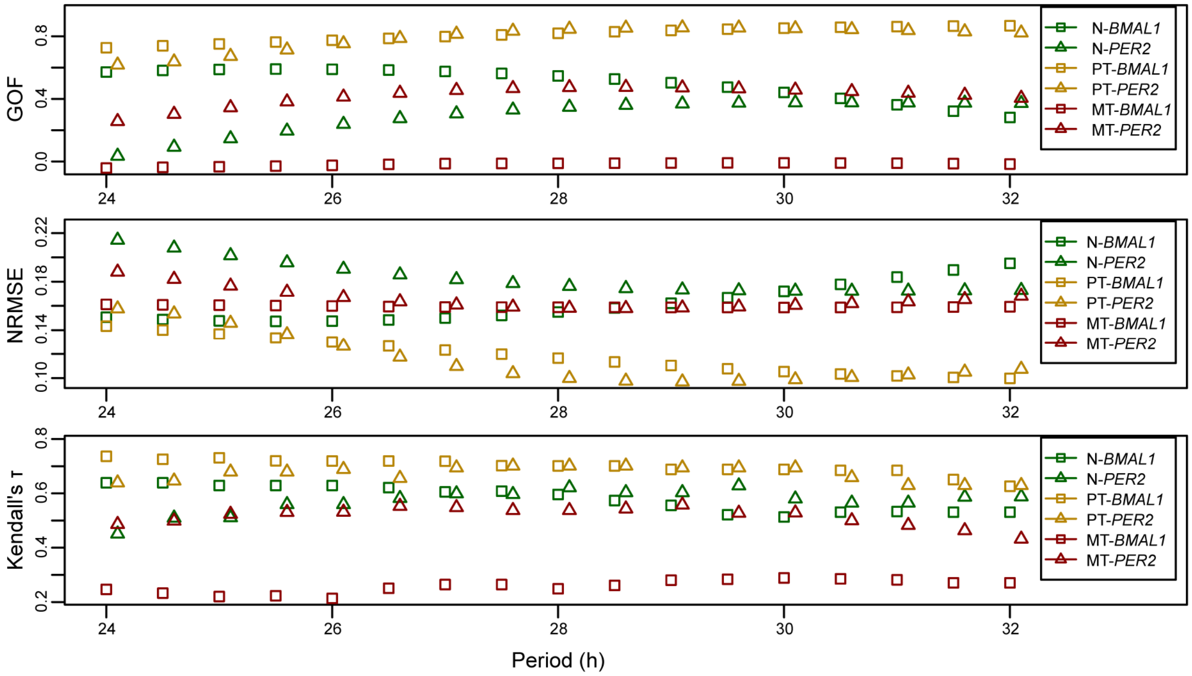


**Figure S1.** *BMAL1* and *PER2* mRNA rhythms in 21PT cells more closely resemble damped sinusoids than those of 21MT-1 cells. Damped cosine curves with fixed periods were fit to time-series with N=6 replicates per time point with the exception of 21MT-1, where N=5 for 4 time points. Fits were computed at a range of periods (24 h to 32 h) and the quality of fits were assessed with three measures: **(above)** GOF (goodness of fit) measures the correlation between the fitted line and the data **(middle)** NRSE (normalized root mean squared error) measures the distance between the data and the fitted lines, and **(below)** Kendall's τ measures the concordance between all pairs of points. Regardless of the period and reporter, the error is smaller (and fit is better) for 21PT cells than for 21MT-1 cells. N=H16N2; PT=21PT; MT=21MT-1.

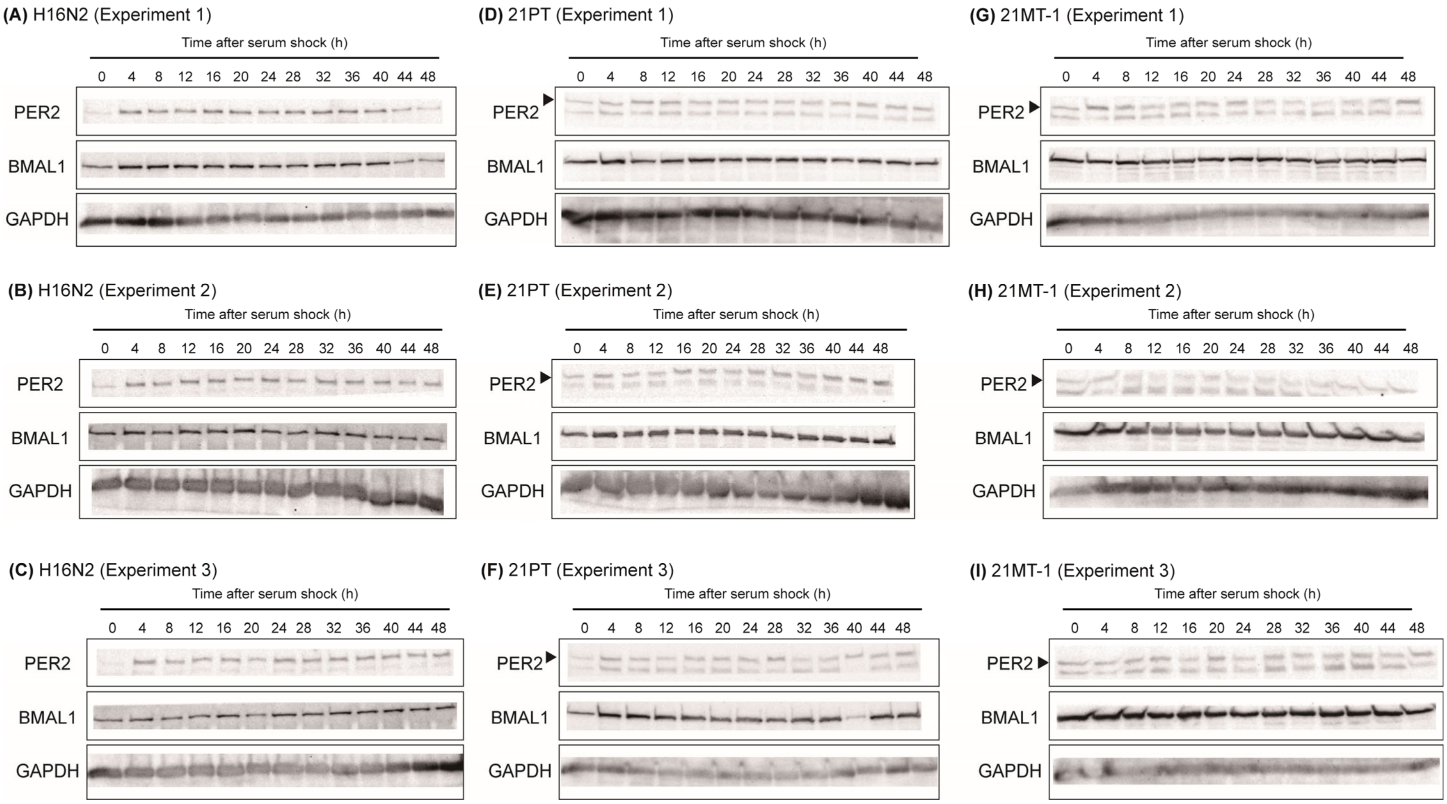


**Figure S2.** All western blots for BMAL1 and PER2 for **(A, B, C)** H16N2 cells, **(D, E, F)** 21PT cells, and **(G, H, I)** 21MT-1 cells.

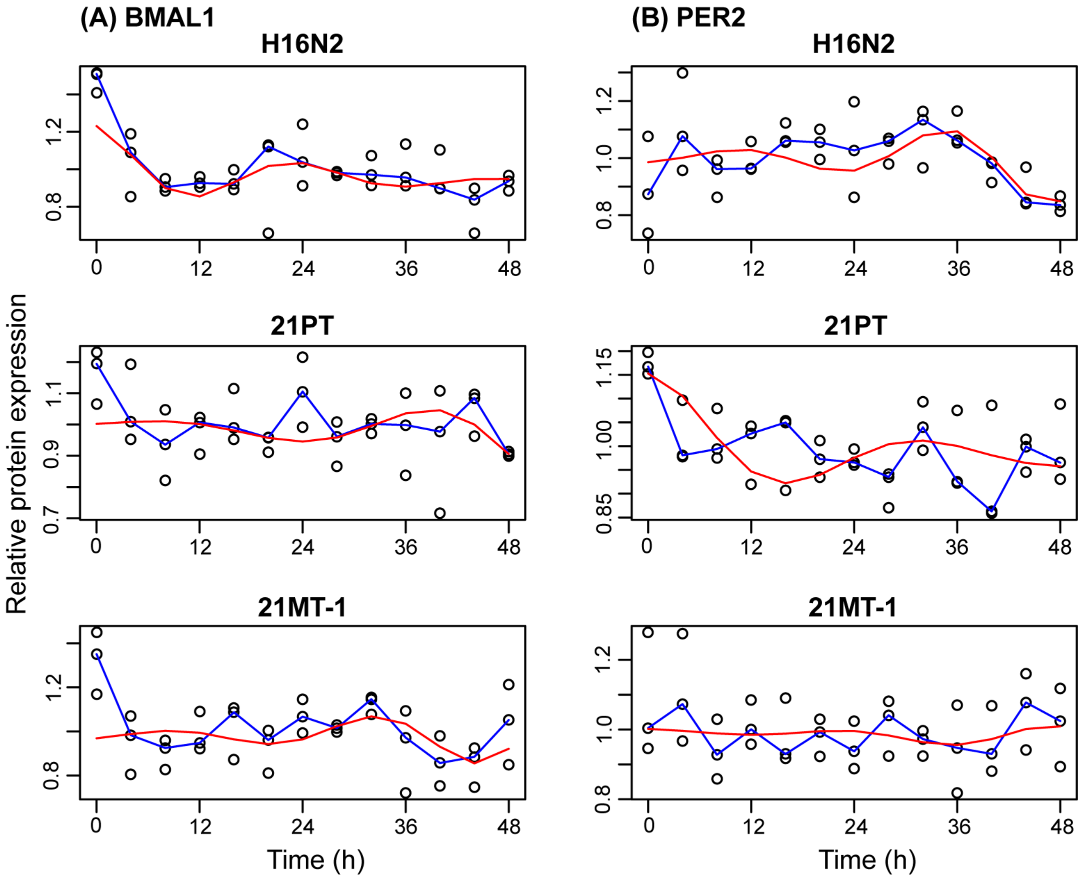


**Figure S3.** Relative protein expression of BMAL1 and PER2 in the 21T series of cells, as determined by western blot. Expression is shown relative to the mean over time. The median signal is shown in blue and the best-fit damped cosine curve is shown in red. Kendall's τ identifies low-quality fits for all time-series (τ=0.28 for BMAL1 and 0.26 for PER2 in H16N2, 0.11 for BMAL1 and 0.20 for PER2 in 21PT, and 0.19 for BMAL1 and 0.18 for PER2 in 21MT-1).

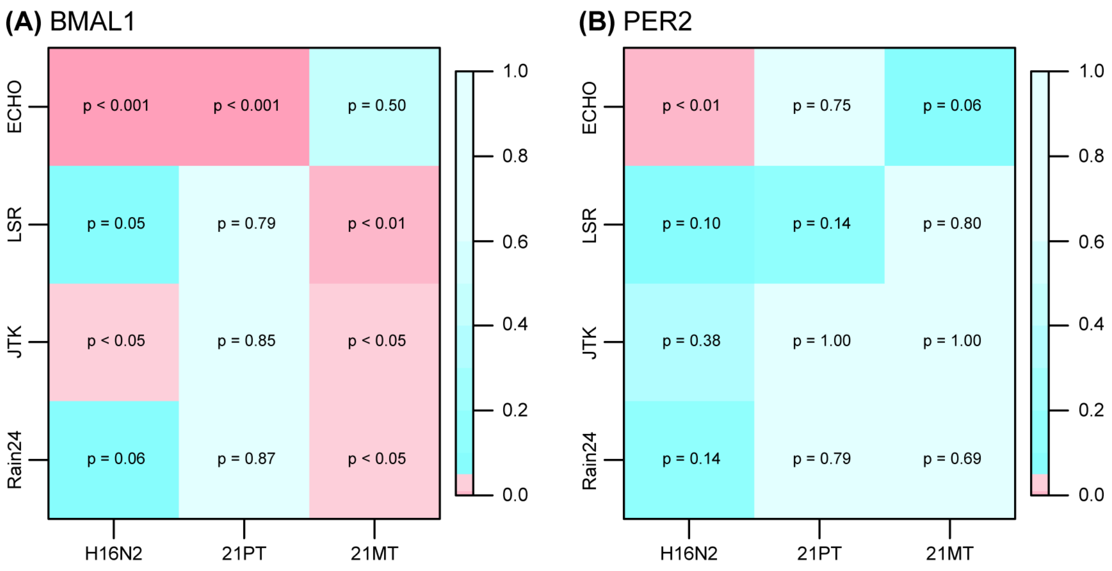


**Figure S4.** P-values for rhythmicity of BMAL1 and PER2 protein expression data. Shown is a heat map with p-values from 4 rhythmicity tests applied to each cell line. There is scant and inconsistent evidence for rhythmicity of protein expression. Rain24 = RAIN with a period of 24 h; JTK = JTK-Cycle; LSR = Lomb-Scargle Permutation test. Values are both labeled and color-coded (shades of red for $p < 0.05$, shades of blue for $0.05 \leqq p \leqq 1$). All tests were applied to time-series with 6 replicates.

**Table S3.** P-values from rhythmicity tests for the BMAL1 protein time-series.

| | **Rain20** | **Rain24** | **Rain28** | **H20** | **H22** | **H24** | **H26** | **H28** | **JTK** | **LS** | **LSR** | **ECHO** |
|---|---|---|---|---|---|---|---|---|---|---|---|---|
| **N** | 0.601 | 0.063 | 1.22e-04 | 0.104 | 0.290 | 0.110 | 1.24e-02 | 2.62e-03 | 2.72e-02 | 0.182 | 0.054 | 9.94e-06 |
| **PT** | 0.060 | 0.874 | 0.415 | 0.170 | 0.206 | 0.287 | 0.259 | 0.224 | 0.852 | 0.999 | 0.788 | 7.52e-05 |
| **MT** | 0.758 | 1.21e-02 | 2.40e-03 | 0.767 | 0.614 | 0.079 | 9.21e-03 | 2.94e-03 | 3.24e-02 | 0.200 | 3.60e-03 | 0.499 |

For each cell line, each time-series with N=3 replicates per time point was tested for rhythmicity, with each of 12 tests. Rain20, Rain24, and Rain28 indicate RAIN with test periods of 20 h, 24 h, and 28h, respectively; H20, H22, H24, H26, and H28 indicate the Harmonic Regression test with periods of 20 h, 22 h, 24 h, 26 h, and 28 h; JTK indicates JTK-Cycle; LS indicates Lomb-Scargle as implemented by MetaCycle; LSR indicates the Lomb-Scargle Permutation test. N=H16N2; PT=21PT; MT=21MT-1

**Table S4.** P-values from rhythmicity tests for the PER2 protein time-series.

| | Rain20 | Rain24 | Rain28 | H20 | H22 | H24 | H26 | H28 | JTK | LS | LSR | ECHO |
|---|---|---|---|---|---|---|---|---|---|---|---|---|
| N | 3.67e-02 | 0.139 | 0.176 | 0.208 | 0.154 | 0.126 | 0.148 | 0.222 | 0.379 | 0.996 | 0.099 | 6.59e-03 |
| PT | 0.847 | 0.793 | 0.471 | 0.646 | 0.820 | 0.512 | 0.315 | 0.279 | 1.000 | 1.000 | 0.142 | 0.752 |
| MT | 0.884 | 0.693 | 0.862 | 0.430 | 0.255 | 0.296 | 0.450 | 0.694 | 1.000 | 1.000 | 0.795 | 0.064 |

For each cell line, each time-series with N=3 replicates per time point was tested for rhythmicity, with each of 12 tests. Rain20, Rain24, and Rain28 indicate RAIN with test periods of 20 h, 24 h, and 28h, respectively; H20, H22, H24, H26, H28 indicate the Harmonic Regression test with periods of 20 h, 22 h, 24 h, 26 h, and 28 h; JTK indicates JTK-Cycle; LS indicates Lomb-Scargle as implemented by MetaCycle; LSR indicates the Lomb-Scargle Permutation test. N=H16N2; PT=21PT; MT=21MT-1.
